# An AI-agent-orchestrated grey-box Transformer framework for sparse pharmacokinetic curve reconstruction and pharmacometric model initialization

**DOI:** 10.64898/2026.05.23.727373

**Authors:** Jingcheng Chen, Jiacheng Wang, Songrui Du, Yangsheng Chen, Kai Li, Jian Song, Dongyang Liu

## Abstract

Clinical pharmacokinetic (PK) modelling is constrained by sparse sampling, limited generalisability of single-drug models, and labour-intensive workflows, making it difficult to infer complete drug exposure from limited concentration observations. We present the Pharmacokinetic Foundation Model (PKFM), a grey-box Transformer framework pre-trained across 32 drugs that reconstructs concentration-time profiles from sparse concentration observations, dosing events, molecular descriptors, and physiological covariates while preserving output interpretability. In representative oral PK curves, three sparse input points recovered the principal absorption-elimination trajectory, achieving coefficient of determination (*R*^2^) = 0.992 for Midazolam oral and *R*^2^ = 0.990 for Verapamil oral. Using reconstructed curves in NONMEM (nonlinear mixed-effects modelling) improved covariance stability and individual prediction accuracy. Contrastive-learning embeddings supported Top-10 physiologically based pharmacokinetic (PBPK) candidate retrieval, with 75.6% of observations within the 2-fold range. A pharmacometrics-informed AI Agent (PM Agent) outperformed general-purpose programming tools in stability and pairwise win rate on a standardised modelling benchmark, with each run requiring human pharmacometrician confirmation before downstream use. These results support cross-drug pre-trained PK models as an information-completion layer for sparse PK evidence and a structured scaffold for the modelling workflow; clinical or regulatory use requires prospective validation, broader external benchmarking, and independent expert assessment.

## 1 Introduction

The central task of quantitative pharmacology is to infer complete in-vivo drug kinetics from limited clinical pharmacokinetic observations, thereby supporting dose optimisation, drug-drug interaction prediction, and extrapolation to special populations[1, 2]. This task has broad applications in drug development, therapeutic drug monitoring, and regulatory evaluation[3], yet its efficiency and quality remain limited by data sparsity, poor model generalisability, and insufficient workflow automation.

Pharmacokinetic (PK) data are inherently sparse, and conventional compartmental models struggle to recover full concentration-time profiles from limited sampling points[4, 5]. Physiologically based pharmacokinetic (PBPK) models offer mechanistic interpretability but span a vast parameter-combination space; identifying the correct combination from sparse data constitutes a high-dimensional inverse problem that is inefficient to solve manually[6, 7]. Existing population pharmacokinetics (PopPK) and PBPK models are typically built fit-for-purpose, limiting cross-drug generalisation and requiring de novo modelling for new compounds or populations[8, 9]. Data-driven artificial intelligence (AI) methods can fit curves but operate as black boxes with insufficient interpretability for regulatory review[10, 11]. Moreover, the modelling workflow relies heavily on expert manual operations and remains non-standardised and inefficient[3, 12]. A useful computational layer should learn shared PK dynamics across drugs, generate auditable outputs from sparse observations, and connect with established modelling workflows.

Here we present the Pharmacokinetic Foundation Model (PKFM) as a grey-box Transformer framework to address these challenges. The framework comprises a reconstruction model and an AI-agent orchestration layer. First, PKFM uses a Transformer encoder-decoder model[13] trained on PBPK-simulated data together with real clinical pharmacokinetic data and physiological representations, learning cross-drug concentration-time sequence embeddings. The model receives sparse concentration observations[14], drug descriptors, and individual physiological parameters, and outputs full concentration-time curve reconstructions; simultaneously, encoder embeddings optimised via contrastive learning[15, 16] support PBPK candidate retrieval, ranking Top-N physiological parameter candidates from a combination library by similarity—each candidate contains complete physiological information (organ volumes, blood flows, body weight, etc.) amenable to expert item-by-item review. Second, it includes a pharmacometrics-informed AI Agent system (PM Agent)[17] that orchestrates a standardised modelling workflow from data preprocessing, curve reconstruction, and structural selection through parameter estimation and diagnostic evaluation, with every decision traceable.

We evaluate PKFM along a sparse-reconstruction-to-pharmacometric-utility chain. The model reconstructs dense pharmacokinetic trajectories from sparse concentration observations across multiple metabolic pathways and routes of administration. Drawing on the principles of symbolic regression[18, 19], we generate compartmental model structure candidates and initial parameter estimates from the reconstructed dense curves; using these as starting points in NONMEM (nonlinear mixed-effects modelling) population pharmacokinetic modelling yields improved covariance stability and individual prediction accuracy relative to direct use of sparse data. The embedding-based retrieval system ranks physiological parameter candidates from the PBPK parameter-combination library for manual review. The pharmacometrics-informed AI Agent (PM Agent) outperforms general-purpose programming agents on a standardised quantitative pharmacology modelling benchmark, with each modelling step requiring human confirmation before progression.

## 2 Results

### 2.1 PKFM links sparse pharmacokinetic evidence to auditable modelling tasks

The PKFM framework converts sparse clinical PK observations into structured pharmacokinetic inferences. Figure 1a shows the human-supervised AI Agent orchestration layer, whose 11 stages cover literature retrieval, data curation, model selection, parameter estimation, diagnostic evaluation and report review; each decision is bound to its input evidence, forming a traceable workflow record. The orchestration layer does not directly participate in concentration prediction but dispatches downstream model outputs and enforces quantitative pharmacology rule checks.

**Figure 1:**
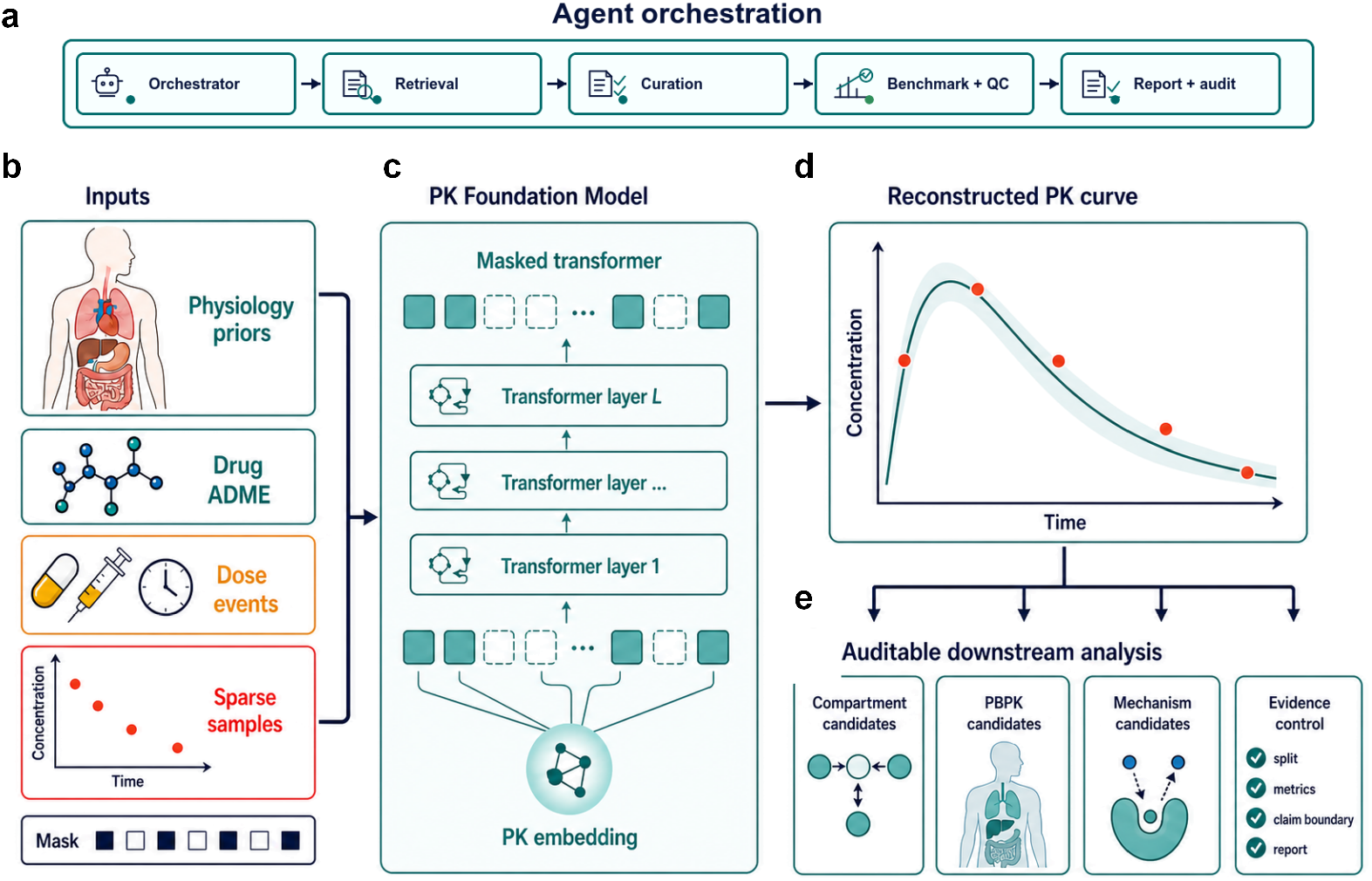
PKFM converts sparse clinical observations into traceable pharmacokinetic inferences. **a**, Human-supervised AI Agent orchestration layer coordinating 11 workflow stages. **b**, Five input categories: drug descriptors, dosing events, sparse observations, physiological priors and availability masks. **c**, Grey-box Transformer core: Transformer encoder–decoder with PK kernel, bounded residual and uncertainty outputs. **d**, Concentration–time curve reconstruction from 3–5 sparse samples. **e**, Three downstream paths: compartmental structure candidates, PBPK candidate retrieval and reviewable mechanistic interpretation.

Figure 1b illustrates the five input categories and their encoding. Drug descriptors carry physicochemical properties and absorption, distribution, metabolism, and excretion (ADME) attributes; dosing events encode dose, timing, route and infusion rate; sparse concentration observations enter the model in the log domain; physiological priors include organ volumes, blood flows, body weight and other individual-level parameters. The feature availability mask flags whether each field is missing, enabling the model to operate under the extensive missingness common in clinical practice without requiring complete inputs.

Figure 1c depicts the Transformer core module of PKFM. The model integrates heterogeneous inputs into a global pharmacokinetic representation and generates concentration predictions at target time points. Predictions are grounded in the multi-exponential decay form of compartmental kinetics, upon which a bounded data-driven correction is applied to accommodate atypical absorption or distribution behaviour, together with point-wise confidence intervals. The model simultaneously learns a pharmacokinetic feature vector for downstream similarity retrieval. This grey-box architecture is designed to keep predictions close to compartmental kinetic shapes and to limit pharmacologically implausible extrapolations from the data-driven correction.

Figure 1d shows the reconstruction process from sparse inputs to dense curves: 3–5 concentration observations enter the model and yield concentration–time predictions spanning the observed time course together with uncertainty bands. Figure 1e presents three traceable downstream analysis paths. Reconstructed curves feed into symbolic regression to produce compartmental model structure candidates and initial parameter estimates. PK embeddings retrieve Top-N PBPK parameter candidates via cosine similarity from a reference library; each combination contains complete organ volumes, blood flows and other physiological parameters available for expert item-by-item review. Mechanistic hypotheses are output as candidates, not as definitive conclusions. The literature PK image extraction pipeline is described in Extended Data Fig. 1.

### 2.2 Sparse clinical samples anchor concentration–time reconstruction

Figure 2 shows in selected clinical PK curves from the Open Systems Pharmacology (OSP) library that when clinical PK curves retain only three concentration points—one each from the early, middle and late phases—PKFM can estimate concentrations at unobserved time points. Figure 2a summarises the model framework; in the OSP curve evaluation, the model uses three-point concentration observations together with drug-name-matched drug descriptors, while dosing information and covariates enter as placeholders. This analysis draws on 476 valid curves from 13 drug–route combinations within the 1,131 records of the Open Systems Pharmacology (OSP) clinical PK library; observations not used as inputs serve for reconstruction evaluation.

**Figure 2:**
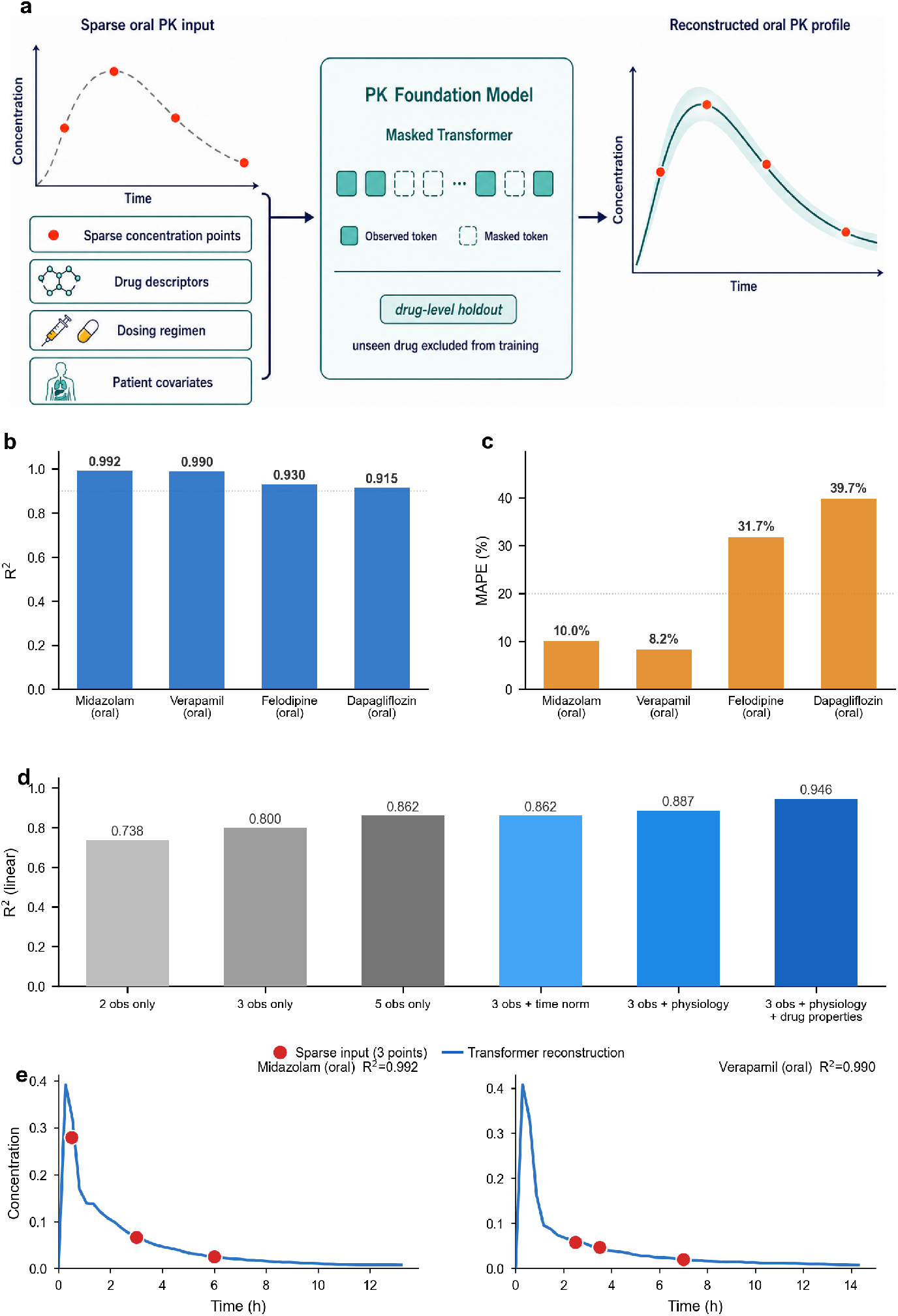
Oral concentration–time curve reconstruction from three-point clinical PK observations. **a**, Model framework schematic. PKFM takes sparse concentration observations as input and can jointly leverage drug descriptors, dosing information and covariates to estimate concentrations at unobserved time points. **b**,**c**, Open Systems Pharmacology clinical PK curves retain one observation each from the early, middle and late phases; remaining observations serve as evaluation references; bar charts show one representative curve per drug meeting the *R*^2^ *>* 0.90 and MAPE*<* 40% selection criteria. **d**, Input ablation on 500 adult core sparse-view training records, comparing contributions of observation count, physiological parameters and drug descriptors to linear-domain *R*^2^. **e**, Two representative reconstruction cases: red dots denote three-point inputs; blue lines denote Transformer-reconstructed curves.

On representative curves meeting the selection criteria, three-point inputs preserve the principal shape information of oral PK curves. Midazolam oral and Verapamil oral achieve coefficient of determination (*R*^2^) = 0.992, mean absolute percentage error (MAPE)=10.0% and *R*^2^ = 0.990, MAPE=8.2%, respectively; Felodipine oral and Dapagliflozin oral yield *R*^2^ of 0.930 and 0.915 with MAPE of 31.7% and 39.7% (Fig. 2b,c). The higher MAPE for the latter two drugs indicates residual concentration errors; the high *R*^2^ values reflect temporal profile concordance between predicted and observed curves.

Figure 2e presents two representative curve examples. For Midazolam oral and Verapamil oral, the three-point inputs do not cover the complete peak shape, yet the model reconstructs the rapid absorption, post-peak decline and terminal elimination phases. These examples illustrate the morphology of the representative high-*R*^2^ curves in Fig. 2b,c and do not independently represent reconstruction accuracy across all OSP curves.

Figure 2d evaluates input information contributions on an independent set of 500 training records with 5-point sparse views. Using concentration points alone, linear-domain *R*^2^ increases from 0.738 at 2 points to 0.800 at 3 points and 0.862 at 5 points; adding physiological parameters raises *R*^2^ to 0.887, and further adding drug descriptors reaches 0.946. These results indicate that both increasing observation count and incorporating contextual information improve *R*^2^ on training records. This ablation does not represent overall error on OSP clinical curves, and MAPE remains 198.9–502.1%.

These results support three-point reconstruction capability on representative oral PK curves and indicate that observation count and contextual information influence *R*^2^ on training records.

### 2.3 Reconstructed curves improve NONMEM estimation quality

Figure 3 places Transformer reconstruction within a population pharmacokinetics (PopPK) modelling context, evaluating how augmenting sparse clinical PK observations into dense concentration– time profiles affects NONMEM estimation quality. The comparison holds the downstream modelling framework constant: both routes use NONMEM First-Order Conditional Estimation with Interaction (FOCE+I); the difference is that the sparse-direct route uses only 3–5 observations per subject, whereas the reconstruction route first augments these to 48–50 time-point concentration–time profiles [4, 20, 21] (Fig. 3a).

**Figure 3:**
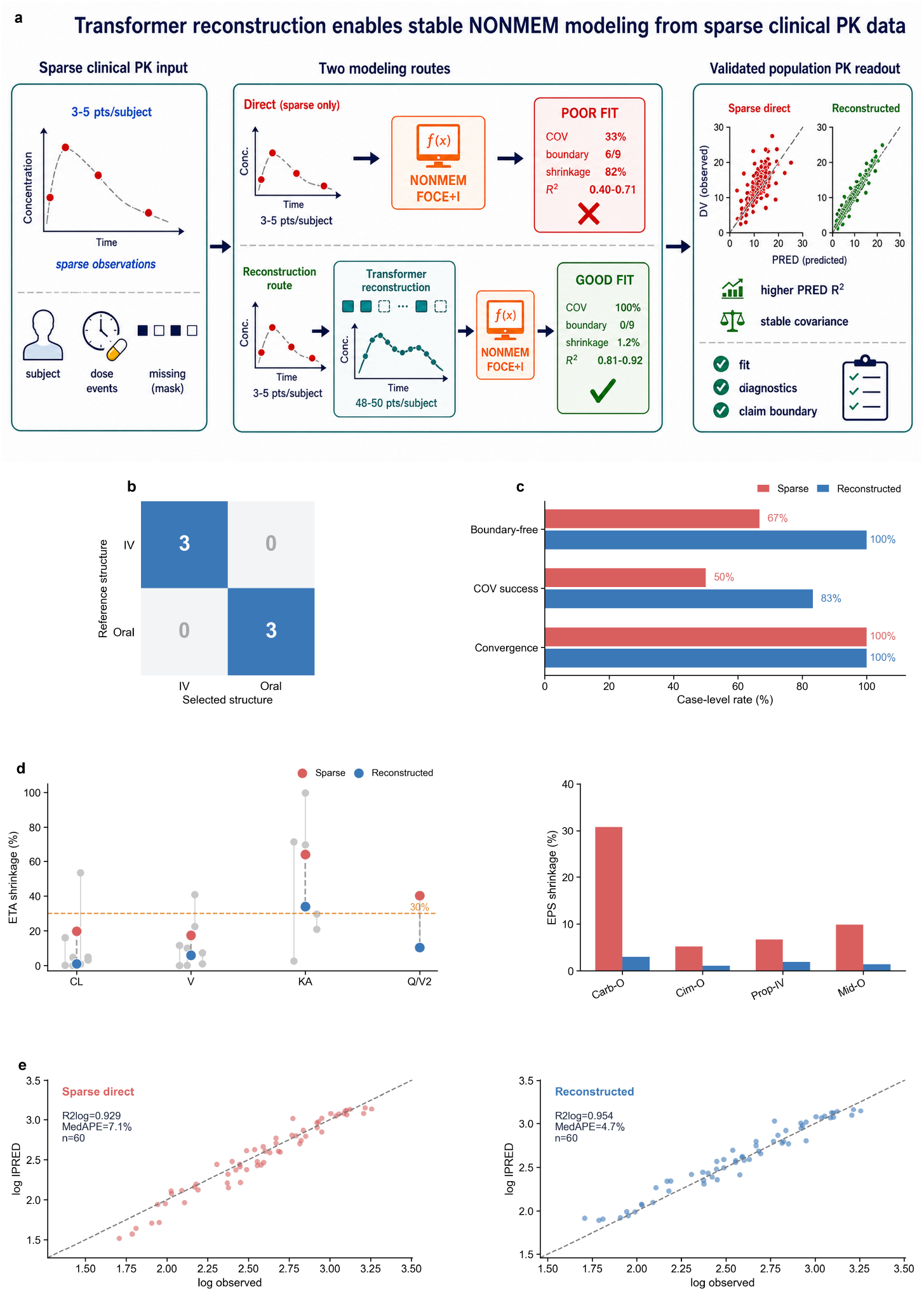
Sparse clinical PK observations reconstructed by the Transformer enter NONMEM FOCE+I modelling for structural selection, estimation diagnostics and individual prediction evaluation. **a**, Schematic of two modelling routes. The sparse-direct route feeds 3–5 concentration observations per subject directly into NONMEM; the reconstruction route first recovers 48–50-point concentration– time profiles via the Transformer before entering the same NONMEM pipeline. Right-side readouts cover prediction concordance, covariance stability, boundary estimates, shrinkage and diagnostic acceptability. **b**, IV/Oral structural selection matrix for six drug–route cases. **c**, Case-level convergence, covariance success and boundary-free estimation proportions for sparse-direct versus reconstruction models. **d**, ETA shrinkage and EPS shrinkage; lower values indicate that inter-individual variability and residual-level information are more fully supported by data. **e**, Observed–IPRED concordance comparison on the same held-out sparse observation set.

Reconstructed curves preserve route-of-administration and structural information. All six drug– route cases are assigned to the correct intravenous (IV) or oral structure, with three IV and three oral cases falling on the diagonal, yielding correct route and structure assignment in this six-case evaluation (Fig. 3b). Curves reconstructed from sparse points retain absorption and elimination morphology sufficient to support structural triage prior to NONMEM entry.

Estimation stability improvements appear primarily in covariance and boundary diagnostics. Both routes achieve 100% convergence, so the comparison focuses on parameter estimation quality. Covariance success rate increases from 50.0% in the sparse-direct model to 83.3% in the reconstruction model, and the boundary-free estimation proportion rises from 66.7% to 100% (Fig. 3c). These results indicate that reconstructed curves provide more continuous temporal information to FOCE+I, facilitating interpretable variance–covariance structures and non-boundary parameter estimates[20, 21].

Shrinkage diagnostics further support this estimation quality improvement. After reconstruction, mean inter-individual variability random effect (ETA) shrinkage on clearance (CL), volume (V), absorption rate constant (KA) and intercompartmental clearance/central volume (Q/V2) decreases from 19.8%, 17.5%, 63.9% and 40.3% to 1.1%, 5.9%, 33.9% and 10.4%, respectively; residual random effect (EPS) shrinkage decreases across all displayed cases, including Carb-O from 30.8% to 3.0%, Cim-O from 5.2% to 1.1%, Prop-IV from 6.7% to 1.9%, and Mid-O from 9.9% to 1.4% (Fig. 3d). Lower ETA and EPS shrinkage indicate that inter-individual variability and residual-level information derive more from observed data, with correspondingly reduced model reliance on population means and error structures[22].

Finally, the reconstruction model yields more accurate individual predictions on the same held-out sparse observation set. The sparse-direct model achieves 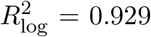 and MedAPE of 7.1%;the reconstruction model improves to 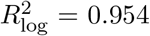 and MedAPE of 4.7% (*n* = 60; Fig. 3e). With evaluation data held constant, the improved prediction concordance and reduced absolute percentage error together indicate that temporal information from Transformer reconstruction propagates through NONMEM parameter estimation and improves subject-level individual predicted concentration (IPRED) readouts.

### 2.4 PBPK candidate retrieval from reference cases

Sparse clinical PK curves are generally insufficient to uniquely determine a set of PBPK parameters[26]. A small number of concentration points capture only local shape features of the exposure–time profile, while dosing information constrains input conditions. This challenge corresponds to the fundamental structure of PBPK models: concentration–time curves are jointly determined by drug-specific parameters, physiological information and dosing conditions[6, 23]. PKFM is trained on PBPK reference curves and their various sparse-sampled versions, using contrastive learning[16] to bring different sparse-sampled versions of the same PBPK parameter combination closer in the representation space[15, 27]. Given a new sparse clinical PK curve, the model therefore performs similarity-based Top-N candidate ranking among validated same-drug PBPK parameter combinations via cosine similarity rather than unique PBPK parameter recovery.

Figure 4 uses 36 Open Systems Pharmacology[28] clinical PK curves covering Alfentanil IV, Verapamil IV, Midazolam IV and Midazolam oral. Each curve is compared against 100 same-drug PBPK parameter combinations (Fig. 4a). Among the top-ranked PBPK parameter combinations, 59.7% of observations fall within the 2-fold range and 25.2% within the 0.8–1.25 range. Expanding evaluation to the top 10 combinations raises these proportions to 75.6% and 42.5%, compared with 51.0% and 17.4% for random parameter combinations (Fig. 4b).

**Figure 4:**
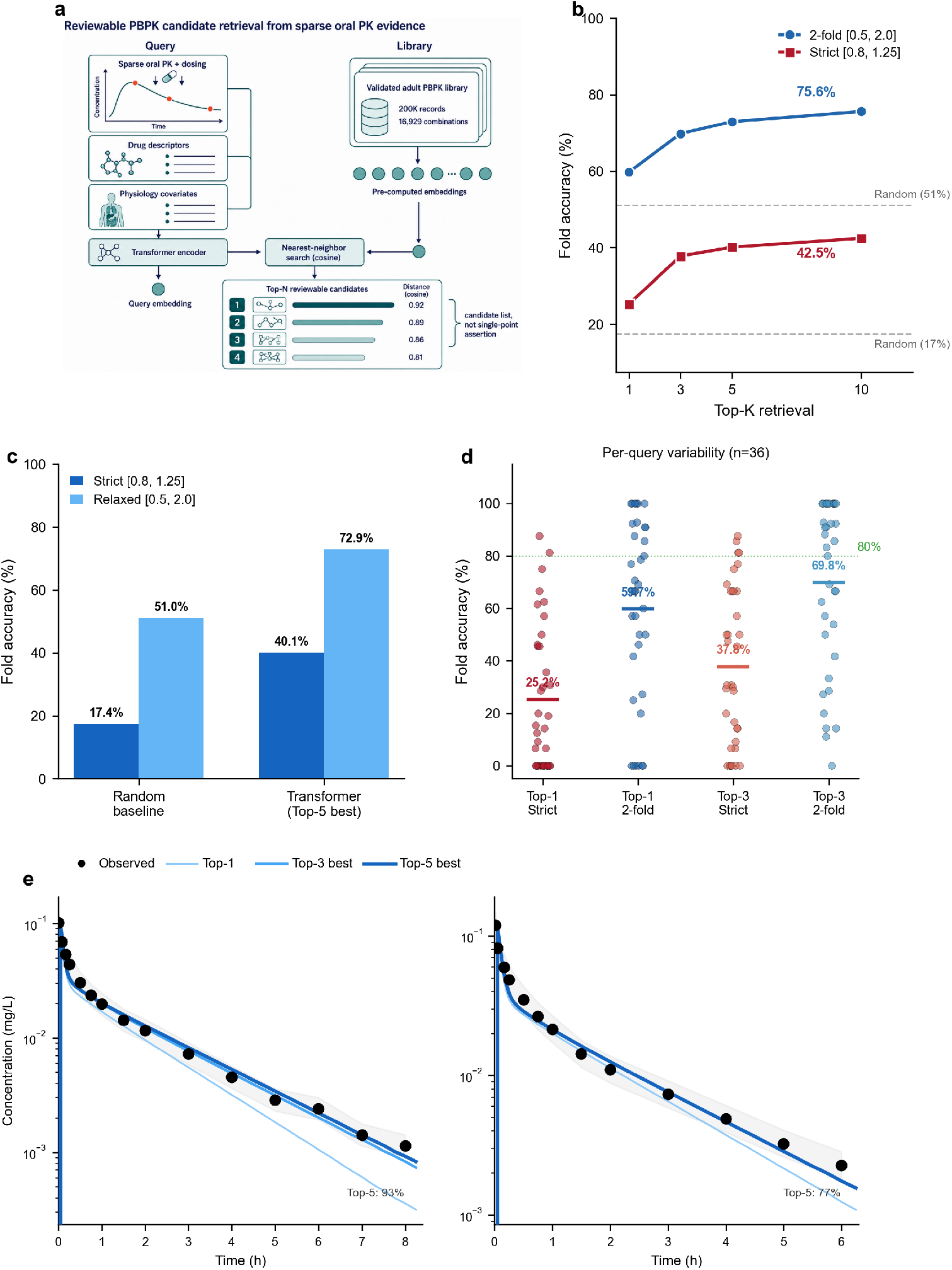
Ranking validated PBPK parameter combinations from real sparse clinical PK observations. **a**, A clinical PK curve under evaluation is characterised by a small number of concentration observations and dosing information. The model encodes this information into a representation vector and ranks same-drug PBPK parameter combinations. Each parameter combination corresponds to a PBPK-simulated concentration–time curve[6, 23]. **b**, Top-K results across 36 real clinical PK curves. For each curve, the best-performing combination among the top K is selected by predicted/observed ratio, and the proportion of observations falling within the 2-fold range[24] and 0.8–1.25 range[25] is reported. **c**, Top-5 results compared with random PBPK parameter combinations. **d**, Top-1 and Top-3 results across different clinical curves. **e**, Two Alfentanil IV cases: black dots denote clinical observations; blue lines denote representative PBPK-simulated curves from the ranked results.

Top-5 provides a scope more amenable to manual review. For each clinical curve, selecting the best-performing combination among the top 5 by predicted/observed ratio yields 72.9% of observations within the 2-fold range and 40.1% within the 0.8–1.25 range. The 2-fold range is a widely used criterion for PBPK prediction accuracy[24], while the 0.8–1.25 range corresponds to the acceptable interval in bioequivalence assessment[25]. Expanding from Top-5 to Top-10 increases the 2-fold and strict-range results by only 2.7 and 2.4 percentage points, respectively (Fig. 4c). This result identifies a small number of parameter combinations most worthy of inspection for continued manual review.

Substantial variability persists across clinical curves. Among Top-3 results, 69.8% of observations fall within the 2-fold range and 37.8% within the 0.8–1.25 range, but individual curve results range from near 0% to 100% (Fig. 4d). In two Alfentanil IV cases, Top-5 results reach 93% and 77%, with corresponding PBPK-simulated curves approximating the rapid elimination phase observed clinically (Fig. 4e).

### 2.5 Pharmacometrics-informed AI Agent (PM Agent) improves modelling bench-mark stability

We compare the human-supervised PM Agent (pharmacometrics-informed AI Agent), Claude Code (Claude Sonnet 4.5), Codex (GPT-5.4) and a general-purpose direct-mean baseline across 27 pharmacokinetic/pharmacodynamic (PK/PD) modelling datasets derived from real clinical modelling cases and 11 standardised workflow stages. The 11 stages follow the modelling process elements specified in the U.S. Food and Drug Administration (FDA) population pharmacokinetics guidance[29] and the International Council for Harmonisation (ICH) M15 model-informed drug development guideline[1], covering data preparation, structural model selection, statistical model specification, covariate screening, parameter estimation, model diagnostics, model validation, sensitivity analysis, simulation design, report writing and regulatory submission preparation. This experiment evaluates whether quantitative pharmacology rules and process checks improve workflow consistency when PKFM curve or representation outputs enter practical modelling pipelines, rather than autonomous clinical or regulatory decision-making. Recent work demonstrates that AI agents can execute end-to-end scientific research workflows under defined task structures[30], whereas general-purpose agent performance in workflows requiring domain-specific judgement such as quantitative pharmacology has not been systematically assessed.

In this AI-rater benchmark, the human-supervised PM Agent achieved the highest total score with the lowest variability. Its mean total score is 63.5 with a median of 64.3, exceeding Claude Code (56.8/58.6) and Codex (52.3/55.1); its standard deviation of 3.6 is lower than 9.3 for Claude Code and 9.4 for Codex (Fig. 5a). Pairwise comparisons are consistent: PM Agent achieves 88.9% win rate against both Codex and the general-purpose direct-mean baseline, and 70.4% against Claude Code (Fig. 5c). These results support pharmacometrics-informed AI-agent processes as a stabilising layer, while human pharmacometrician confirmation remains necessary at each modelling step.

**Figure 5:**
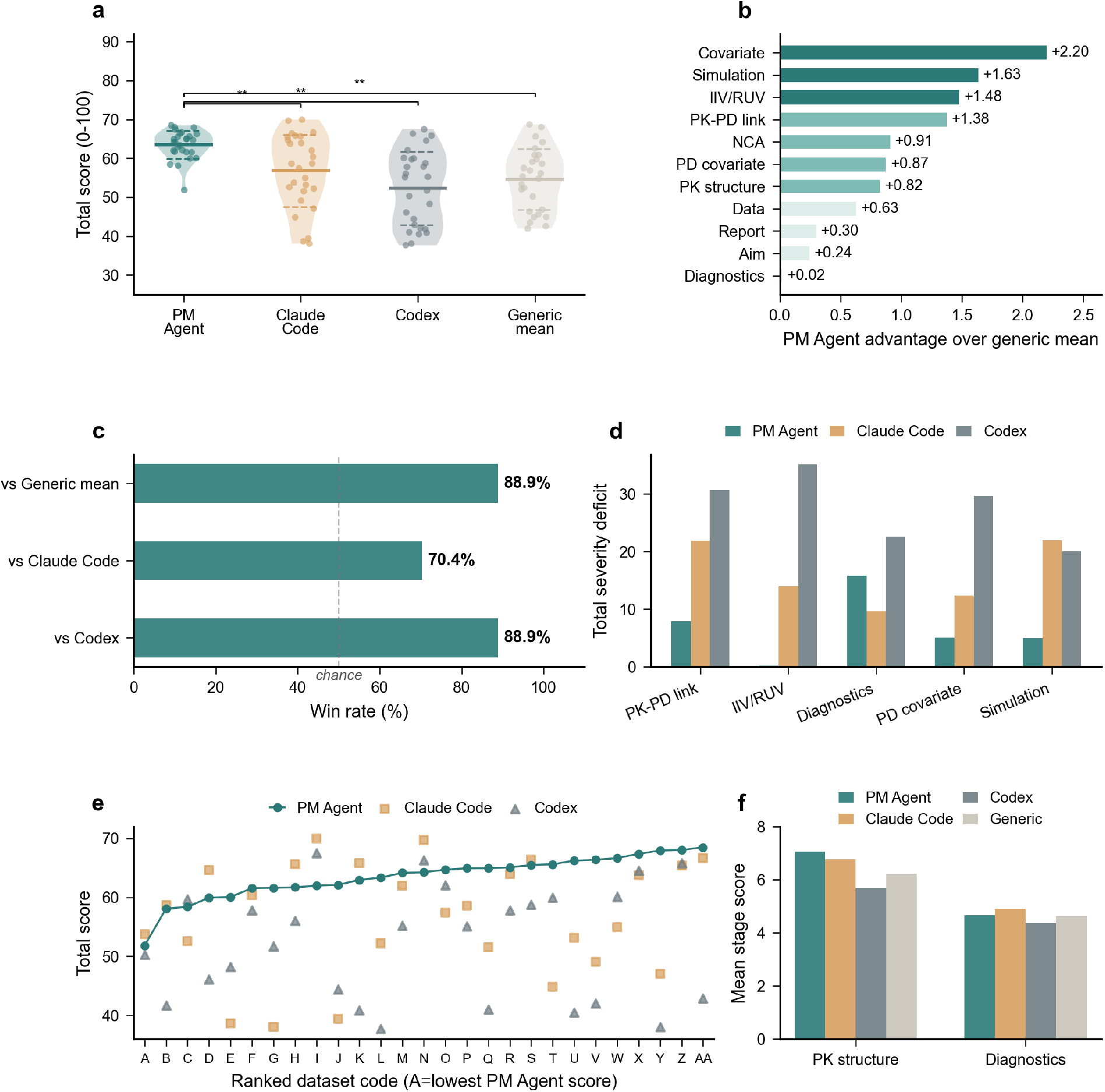
Benchmarking the human-supervised pharmacometrics-informed PM Agent against general-purpose programming agents on quantitative pharmacology modelling workflows. **a**, Total score distributions across 27 PK/PD modelling datasets. **b**, Stage-level advantages of PM Agent relative to the general-purpose direct-mean baseline. **c**, Pairwise win rates of PM Agent against Codex, Claude Code and the general-purpose direct-mean baseline. **d–f**, High-severity gap stages, dataset trajectories sorted by PM Agent total score from low to high, and stage scores for PK structural selection and diagnostic evaluation.

Advantages concentrate in modelling-judgement-intensive stages. The largest stage-level gains of PM Agent over the general-purpose direct-mean baseline appear in covariate model selection (+2.20), simulation design (+1.63), inter-individual variability and residual error model specification (+1.48), and PK–PD linkage (+1.38) (Fig. 5b). These stages require simultaneous consideration of drug exposure, variability sources, model interpretability and simulation objectives, aligning with the critical decision nodes emphasised in published population modelling best-practice guidelines[31] and the Model-Informed Drug Discovery and Development (MID3) framework[2]; the results support the value of quantitative pharmacology rules in complex modelling decisions.

The benchmark also identifies remaining weak points. In PK structural selection, PM Agent achieves a mean stage score of 7.06, exceeding Claude Code, Codex and the general-purpose mean baseline; in diagnostic evaluation, PM Agent scores 4.67, comparable to the general-purpose mean baseline and below Claude Code at 4.92 (Fig. 5f). Figure 5 supports improved workflow reliability, particularly in structural, covariate, variability model, PK–PD linkage and simulation design stages; diagnostic interpretation and robustness judgement still require human review.

## 3 Discussion

PKFM evaluates the feasibility of a computational framework that maps sparse PK observations to auditable quantitative pharmacology outputs through cross-drug pretrained generalisation—a step beyond purely per-drug AI-PK approaches. Neural ordinary differential equations (ODEs)[32, 33], transfer learning, and Transformer longitudinal analysis[34] have improved data efficiency but remain within per-drug paradigms; general-purpose time-series foundation models[35, 36] generalise across domains but lack PK physical constraints. PKFM addresses both limitations through joint pretraining on PBPK simulations and real data, providing preliminary evidence for transfer across selected held-out drugs[37]. The grey-box architecture embeds auditability as a native constraint[38, 39, 40]: outputs follow compartmental multi-exponential decay, bounded residual corrections are capped at threefold, and the model degrades to mechanistic baseline predictions when evidence is insufficient. For PBPK parameter estimation, where Bayesian sampling is computationally expensive and machine learning (ML) surrogates yield point estimates without auditable candidate spaces[41, 42, 43], PKFM uses a retrieval strategy that narrows candidates to a ranked shortlist for physiological plausibility review.

Current evidence supports framework feasibility but explicit gaps remain before clinical application. Thirty-two drugs are insufficient to represent clinical diversity; publicly available OSP data do not constitute prospective clinical validation[44]; individual physiological parameters are typically incomplete in practice; and agent evaluation employed AI raters rather than human experts. From proof of concept to clinical adoption, systematic external validation, explicit drug-level and route-level held-out testing, calibration analysis, and human expert evaluation are required[45].

The long-term value of PKFM lies in bridging PopPK and PBPK workflows. Recent automated modelling tools—pyDarwin[46], NSGA-II/III[47], ML-assisted selection[48], and nlmixr2auto[49]— address model selection automation but assume sufficient data information content. PKFM addresses the upstream problem of supplementing temporal profiles when sparse data cannot support reliable estimation; the two are complementary. Near-term priorities include expanding drug coverage and therapeutic drug monitoring (TDM) validation, medium-term integration of pharmacogenomics for special populations, and reproducible evaluation packages for model-informed drug development under the ICH M15 framework[38, 1]. Multi-agent systems have been explored for scientific discovery in chemistry and biomedicine[50, 17]; the PKFM agent layer should remain a human-supervised AI Agent system, with each run reviewed before clinical or regulatory use, until stronger validation is available.

## 4 Methods

### 4.1 Training data generation

Individualised pharmacokinetic prediction in clinical studies faces two fundamental data bottlenecks. Real clinical data, constrained by ethics approvals and sampling density, cannot cover a sufficient drug–population–physiological-state combinatorial space; relying solely on simulated data may introduce systematic bias. To resolve this tension, we adopted a dual-source training strategy combining PBPK mechanistic simulation with real literature data extraction, enabling the model to learn both mechanism-driven physiology–PK mapping relationships and the distributional characteristics of real clinical data.

PBPK simulation data were generated using PK-Sim 12.3.51 on the Open Systems Pharmacology platform[51, 28]. We selected 48 adult drug models and 17 pregnancy models from the OSP validated model library, covering 56 drugs. To enable the model to learn the systematic effects of physiological parameter variability on PK behaviour, the simulation design incorporated multi-dimensional population perturbations spanning six ethnic populations (European ICRP, Asian Tanaka, Japanese, Black American, Mexican American, White American) with corresponding body-size and organ-proportion differences, ages from 18 to 90 years, health states including healthy (70%), chronic kidney disease (CKD) (15%) and hepatic impairment (HI) (15%), and four metabolic enzyme phenotypes: poor metaboliser (PM), intermediate metaboliser (IM), normal metaboliser (NM) and ultra-rapid metaboliser (UM). The full-factorial combination of these dimensions yields a total design space of approximately 48 million simulations (961 drug scenarios multiplied by approximately 50,000 virtual individuals); the current training uses the adult core subset. The PK-Sim virtual population generator, based on known physiological and anthropometric parameter distributions[52], produces virtual individuals with realistic physiological variability. Raw simulations yielded 1,114 dense concentration–time curves; after rigorous quality filtering (excluding biologics, retaining only adult small molecules) we obtained 5,649 validated anchor curves covering 32 drugs. Each curve is accompanied by 96-dimensional physiological system parameters encompassing organ volumes, blood flow rates, tissue composition fractions, glomerular filtration rate and gastrointestinal transit parameters, extracted directly from the PK-Sim virtual population generator. This simulation strategy preserves the causal chain between physiological parameters and PK curves, enabling the model to learn the correspondence between physiological characteristics and pharmacokinetic behaviour while reducing extrapolation risk inherent in purely statistical fitting.

Real literature data were sourced from approximately 30,000 quantitative pharmacology publications. We developed a PK figure visual extraction system based on YOLOv8[53] object detection and coordinate regression, automatically extracting concentration–time data points from published figures. The system simultaneously extracts patient physiological metadata (body weight, age, renal function, dosing regimen, etc.), achieving matching between patient physiological characteristics and pharmacokinetic data so that literature-extracted data possess the same “physiological parameters– PK curve” paired structure as the PBPK simulation data. After integration, both data types are used jointly for Transformer training: PBPK data provide a broad mechanistic foundation, while literature data provide calibration signals from real clinical distributions.

Clinical PK sampling is typically sparse (2 to 5 time points), and patient information is often incomplete. To adapt the model to these real-world constraints, we generated multiple sparse training instances from each dense anchor curve, ultimately producing 200,000 training instances (168,395 training, 31,605 validation). Sparse sampling covers six clinically common patterns: two-point, three-point, five-point, missing absorption phase, missing elimination phase and trough-only. Data augmentation includes log-domain observation noise injection, sampling time jitter, feature missingness masks and linear dose scaling. The 32 drugs were split at a 4:1 ratio using a drug-level split (26 training, 6 validation)[54, 55], supporting an internal test in which validation drugs are not used for training within this split.

Independent validation data came from two external sources: published clinical study observations curated in the OSP model library (1,131 curves, 23 drugs) and the PK-DB pharmacokinetic database[56] (803 studies, 2,863 curves, 829 publications). Neither dataset participated in model training. The 23 drugs in the OSP validation set partially overlap with the 32 training drugs, so these analyses should be interpreted as external-record evaluation rather than uniformly unseen-drug validation; the curve reconstruction evaluation in Fig. 2 uses 476 valid curves from 13 drug–route combinations in the OSP validation set, where the model receives only sparse observation points as input without access to the remaining observations of each curve.

### 4.2 PKFM grey-box architecture

The central challenge in pharmacokinetic concentration prediction is balancing interpretability with expressive power. Pure mechanistic models (compartmental models) have interpretable parameters but are limited in expressiveness by their pre-specified structure; pure data-driven models (black-box neural networks) are flexible but produce outputs lacking pharmacological meaning. The Transformer module of PKFM resolves this tension through a grey-box architecture[57, 11], combining mechanism-driven pharmacokinetic kernel functions with data-driven neural network residual correction, so that model outputs remain constrained by pharmacokinetic structure while possessing adaptability to complex PK behaviours.

The core prediction equation of the model is

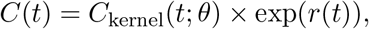

where the kernel function *C*_kernel_ provides a multi-exponential decay form consistent with fundamental pharmacokinetic laws, and the bounded residual term *r*(*t*) compensates for nonlinear effects that the kernel cannot capture (maximum 3-fold correction). The key constraint of this design is the boundedness of the residual:

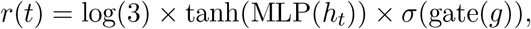

where the confidence gate approaches zero when evidence is insufficient, causing the residual to vanish automatically and the model to degenerate to the pure mechanistic kernel function. When input information is sparse or the model is uncertain, predictions are encouraged to remain close to the pharmacokinetic prior, reducing the risk of uncontrolled extrapolation by black-box models under data scarcity.

The total model parameter count is 22.8M. Input consists of four token sequence types: drug descriptors (12-dimensional, encoding physicochemical properties and metabolic characteristics), dosing events (up to 28, each containing time, dose, rate and route of administration), sparse concentration observations (up to 12 time points) and covariates (192-dimensional: 96-dimensional normalised physiological parameters plus 96-dimensional feature masks indicating parameter availability). Temporal information is processed through continuous time encoding[58]: 64 log-uniformly spaced base frequencies (0.01 to 100) undergo sine–cosine transformation to generate 128-dimensional representations, then linear projection to the 512-dimensional model space, enabling the model to handle arbitrary PK time scales from minutes to days.

The Context Encoder employs a Pre-norm Transformer architecture[59] (4 layers, 4 attention heads, *d*_model_ = 512, *d*_ff_ = 2048, dropout 0.15), performing self-attention encoding over all input tokens and outputting sequence-level context representations and a global context vector. The Query Decoder is a 2-layer, 4-head Cross-attention Transformer[13] that uses time encodings of target prediction time points as queries, extracting information relevant to each prediction instant from the encoder context via cross-attention.

The Kernel Head predicts interpretable PK response kernel parameters from the global context vector. The kernel function is a 4-term multi-exponential decay model:

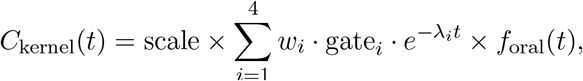

where rate constants *λ*_1_ *< λ*_2_ *< λ*_3_ *< λ*_4_ are guaranteed to be ordered via cumulative softplus, *w*_*i*_ are normalised weights, soft-select gating controls the number of active terms through a learnable softness parameter, and the oral absorption factor 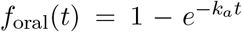is activated by an oral gate. This parameterisation naturally corresponds to the analytical solution forms of 1-to 4-compartmental models[60, 61], enabling the model to adaptively express PK dynamics of varying complexity without pre-specifying compartmental structure. The Uncertainty Head outputs Student-*t* distribution parameters (degrees of freedom *v >* 2.1, heteroscedastic scale *σ*(*t*)) and an out-of-distribution (OOD) detection score, providing uncertainty estimates for each prediction time point[62]. The core advantage of this grey-box design is that kernel function parameters directly correspond to pharmacokinetic concepts (elimination rate, distribution rate, absorption rate), making model predictions directly auditable by pharmacologists. The bounded residual provides adaptability to non-standard PK behaviours (such as nonlinear elimination, enterohepatic circulation) while maintaining interpretability, and the balance between data-driven and mechanism-driven components is automatically regulated by the confidence gate mechanism.

### 4.3 Contrastive learning and retrieval

The encoder of the PKFM Transformer core outputs a global context vector. During pre-training this vector primarily serves concentration prediction, and its geometric structure requires further organisation to support similarity-based retrieval. In clinical applications, rapidly retrieving reference cases with similar physiology–pharmacokinetic characteristics from a historical database given a new patient’s sparse PK observations may support individualised dosing review. The purpose of contrastive learning is to reshape the geometry of the embedding space so that different sparse views of the same drug–population–physiological-parameter combination cluster together in vector space, while views from different combinations are pushed apart.

Positive samples are defined as different sparse sampling views of the same anchor curve (i.e., the same drug, population and physiological parameter combination), reflecting identical underlying PK behaviour with only the observation pattern differing. Negative samples are views from different anchors. Each batch samples 512 anchors with 4 views per anchor, yielding an effective batch size of 2,048. Training uses a supervised contrastive learning (SupCon) variant of the noise-contrastive estimation loss (InfoNCE loss)[16, 27] with temperature parameter *τ* = 0.07. The joint training strategy combines reconstruction loss and contrastive loss with weighting (*ℒ* = 0.3 *× ℒ*_recon_ + 1.0 *× ℒ*_CL_): reconstruction loss maintains concentration prediction capability while contrastive loss optimises embedding retrieval performance. The contrastive learning weight is set to 1.0 and reconstruction weight reduced to 0.3, reflecting that the primary optimisation objective at this stage is structuring the embedding space while preventing degradation of predictive capability. After contrastive learning, the model’s embedding space supports efficient nearest-neighbour retrieval: given a new patient’s sparse observations, the most PK-behaviourally similar candidates can be identified from thousands of reference curves as starting points for downstream parameter review.

### 4.4 Symbolic regression and NONMEM evaluation

The Transformer model reconstructs dense concentration–time curves from sparse observations. Clinical pharmacology practice further requires expressing PK behaviour as specific compartmental model parameters (clearance CL, volume of distribution V, absorption rate constant KA, etc.). Symbolic regression[63], as an independent downstream analysis step, converts the continuous curves output by the model into interpretable compartmental model parameters. Symbolic regression and the model’s internal Kernel Head operate at different levels: the Kernel Head provides the internal grey-box scaffold during prediction generation; symbolic regression post-processes the model’s final output curves and extracts reviewable compartmental parameter candidates.

The specific procedure is as follows: for each Transformer-reconstructed complete concentration– time curve, nonlinear least-squares fitting (`scipy.optimize.curve_fit`[64], log-domain) is performed against pre-defined candidate equation families. The candidate equation families include the 1-compartment intravenous model (mono-exponential decay), 2-compartment intravenous model (bi-exponential decay) and 1-compartment oral model (Bateman function)[60, 61]. The optimal compartmental structure is determined by comparing fitting residuals across candidates, yielding candidate parameter values under that structure. This design bridges the Transformer’s flexible predictive capability with traditional pharmacokinetic parameterised expression: the model is responsible for reconstructing complete curves from sparse data, while symbolic regression converts curves into the parametric language familiar to pharmacologists. With the two tasks decoupled, even when underlying PK behaviour deviates from simple compartmental models, the Transformer can still reconstruct curves while symbolic regression provides the best-approximating parameterised description for expert review.

The NONMEM evaluation compares the sparse-direct route with the reconstruction route. In the sparse-direct route, each subject’s 3–5 clinical PK observations are input directly into NONMEM; in the reconstruction route, the Transformer first completes a 48–50 time-point concentration–time profile, which is then modelled using the same NONMEM workflow. The evaluation covers route-of-administration or structural model selection, convergence and covariance step success, boundary hits, parameter relative standard error (RSE), ETA or EPS shrinkage, and IPRED error on the same sparse dependent variable (DV)[4, 20, 21, 22].

### 4.5 AI Agent orchestration and benchmark

Translating PKFM outputs into an end-to-end modelling workflow requires standardised task orchestration. In traditional quantitative pharmacology practice, the complete pipeline from data quality control through model selection, parameter estimation, diagnostic evaluation and report generation involves extensive expert judgement and manual operations. The human-supervised AI Agent orchestration system[17] is implemented as a high-level OrchestratorAgent linked to a PPKModelingAgent, a tool registry and NONMEM/template/reporting tools, and structures this pipeline into 11 executable stages. Each stage records its inputs and outputs; a human pharmacometrician reviews each run result before the next stage is accepted or used downstream.

We constructed the population pharmacokinetic benchmark (PPK-Bench) to evaluate the modelling capability of Agent systems. This benchmark comprises 27 independent PK/PD modelling datasets, all derived from real clinical modelling cases accumulated by our research group, covering the complete 11 stages of quantitative pharmacological modelling (from problem definition, data quality control, exploratory analysis, PK structural model selection, residual error modelling, covariate screening, diagnostic evaluation, PK/PD linkage through to simulation and reporting), with a total score of 100 allocated across stages. PKFM’s curve reconstruction and parameter retrieval capabilities are integrated at Stage 3 (PK structural model selection), providing the Agent with compartmental structure candidates and initial parameter estimates, replacing the model selection step that traditionally relies on expert experience. Three Agent systems were evaluated under identical conditions: PM Agent (the human-supervised pharmacometrics-informed AI Agent integrating PKFM outputs), Claude Code (based on Claude Sonnet 4.5) and Codex (based on GPT-5.4). Each Agent’s response to each dataset was independently scored by 3 artificial-intelligence (AI) raters (Gemini 3 Flash, DeepSeek V3.2, Grok Code Fast 1), producing 243 scoring records (27 datasets multiplied by 3 Agents multiplied by 3 raters). The multi-rater design reduces dependence on any single rater; the three raters come from different model families, and their inter-rater agreement is reported as a screening-level reliability indicator that should not replace human pharmacometrician evaluation.

### 4.6 Software and computing environment

PBPK simulations used PK-Sim 12.3.51 (Open Systems Pharmacology platform[51]) with the OSPSuite-R 12.1 interface for batch execution. Deep learning used Python 3.10, PyTorch 2.1[65] and CUDA 12.1. Numerical computation used NumPy 1.24 and SciPy 1.11[64]. Data processing used pandas 2.0; visualisation used matplotlib 3.8 and seaborn 0.13. Figure data extraction used YOLOv8 (ultralytics 8.1)[53]. Statistical analysis used R 4.5. Model training was performed on NVIDIA A100 80GB PCIe GPUs; batch PBPK simulations used a 258-node cluster with 64 CPU cores per node.

## 5 Data and code availability

### Data availability

This study used two publicly available pharmacokinetic data sources. Open Systems Pharmacology (OSP) models, simulation resources, and published clinical observation data were obtained from the OSP ecosystem and its model repository[51]. The PK-DB pharmacokinetic database served as the source for literature time-course data and associated metadata[56].

The model input tables, figure source data, benchmark summary tables, and cleaned intermediate datasets generated in this work will be organised as a reproducibility package before journal submission. Because these files contain reprocessed versions of third-party public data, model-training derivatives, and local evaluation artefacts, the release scope will follow original data licences, journal requirements, and reproducibility needs, with source-data provenance, preprocessing scripts, figure source tables, and evaluation outputs identified where possible.

### Code availability

The model training code, data processing scripts, benchmarking tools, and manuscript assembly scripts developed in this study will be prepared as a minimal reproducibility package where licence conditions permit. Should the target journal or peer-review process require code inspection, restricted-access instructions can be provided for components that cannot be publicly released.

## Acknowledgements

This work was supported by High-performance Computing Platform of Peking University.

## Extended Data

**Extended Data Fig. 1 — Literature concentration–time curve extraction trained on pharmacokinetic images**

**Extended Data Fig. 1.**
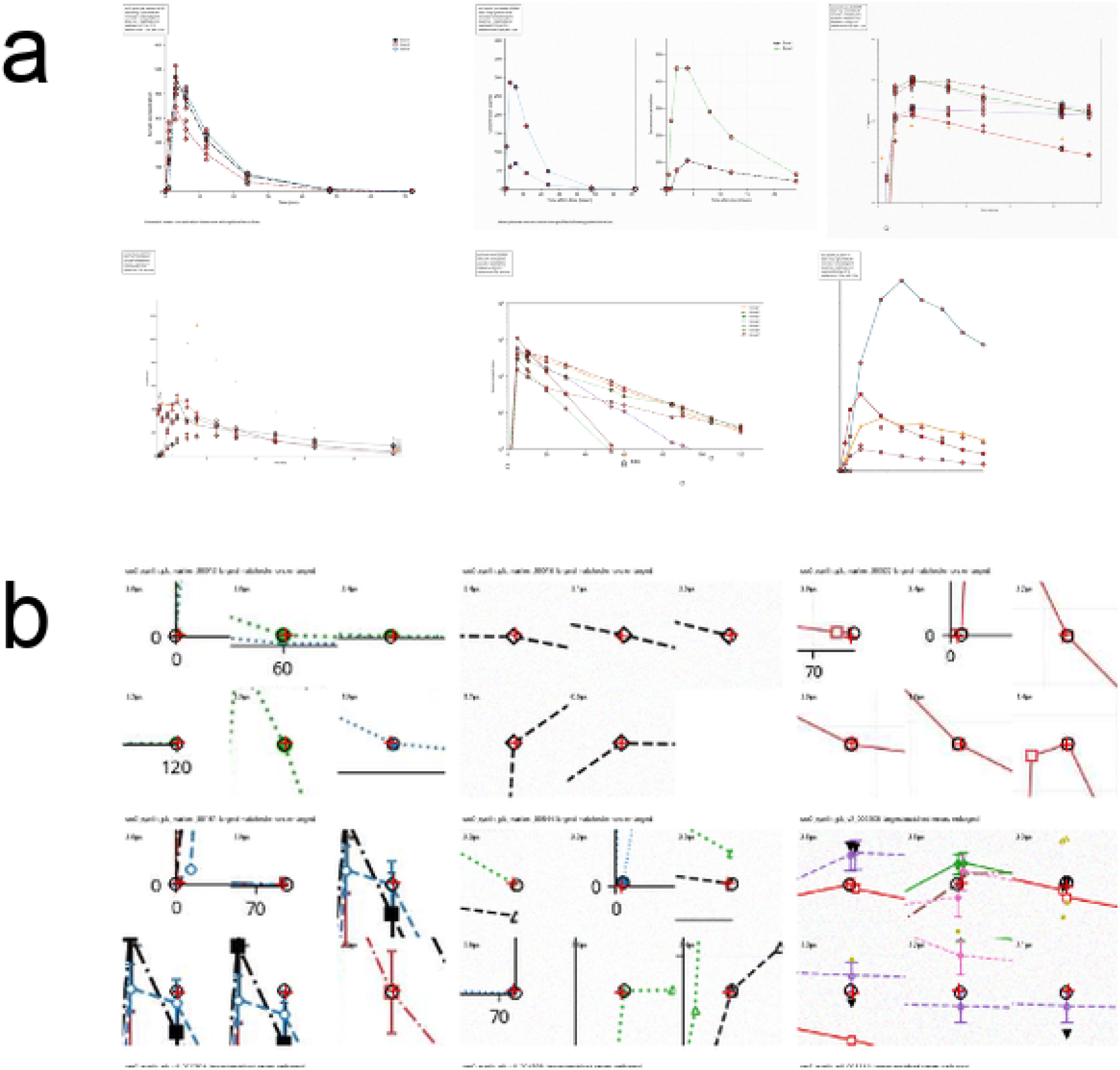
A pharmacokinetic image extraction model trained on PK figure data locates concentration–time data points from literature-style PK curves. **a**, Examples of source-retained PK images used for training and evaluation, covering single-curve and multi-curve plots, linear and logarithmic axes, error bars, sparse sampling and diverse marker styles. **b**, Point localisation results of the model on representative images. Red markers indicate extracted data points, which are subsequently mapped to time–concentration records via axis coordinate transformation.

Literature PK images are an important supplementary source of PKFM training data. Previous PK-DB work has demonstrated that time-course data from pharmacokinetic literature require figure digitisation, unit standardisation and validation rule processing before they can enter a computable database[1]. Accordingly, we trained a pharmacokinetic image-specific extraction model to locate curve data points from literature-style concentration–time figures; the model implementation follows an object detection approach that can be trained on custom data[2].

The training and evaluation sets retain common literature image scenarios including multi-curve plots, logarithmic concentration axes, error bars, sparse sampling and marker overlap. For each image, the original time–concentration values, axis mappings and data-point centre annotations are simultaneously saved, enabling pixel-level localisation error and recovered PK data error to be traced on a per-image basis.

The benchmark comprises 200 PK figures, 4,237 source/gold data points and 3,806 matched points. Point-centre detection precision is 0.9211, recall is 0.8983 and F1 is 0.9095; the median pixel error of matched points is 0.4659 px, median time error is 0.0261 h and median log_10_ concentration error is 0.00134. Cmax, Tmax and partial AUC errors are used to verify whether recovered points preserve peak exposure, time to peak and local exposure information; these metrics are consistent with commonly used parameters for concentration–time curve exposure assessment[5].

The pharmacokinetic image-trained point extraction model converts literature-style PK curves into auditable time–concentration candidate records, providing an image-based entry point for subsequent literature data curation and PKFM training data expansion. These results are limited to the source-retained benchmark and the current image extraction pipeline; real PDF layout parsing, figure-caption metadata matching, unit standardisation and QC-retained model-ready dataset construction require independent validation.

## References

[1] International Council for Harmonisation. M15 General Principles for Model-Informed Drug Development, 2026. ICH Guideline, Step 4, adopted 29 January 2026.

[2] S. F. Marshall, R. Burghaus, V. Cosson, et al. Model-informed drug discovery and development: current industry good practice and regulatory expectations and future perspectives. CPT: Pharmacometrics and Systems Pharmacology, 8(2):87–96, 2019.

[3] R. Madabushi et al. Review: Role of model-informed drug development approaches in the lifecycle of drug development and regulatory decision-making. Pharmaceutical Research, 39(8):1669–1680, 2022.

[4] D. R. Mould and R. N. Upton. Basic concepts in population modeling, simulation, and model-based drug development—part 2: Introduction to pharmacokinetic modeling methods. CPT: Pharmacometrics and Systems Pharmacology, 2(4):e38, 2013.

[5] L. B. Sheiner and S. L. Beal. Evaluation of methods for estimating population pharmacokinetic parameters. i. michaelis-menten model and experimental pharmacokinetic data. Journal of Pharmacokinetics and Biopharmaceutics, 8(6):553–571, 1980.

[6] L. Kuepfer, C. Niederalt, T. Wendl, et al. Applied concepts in pbpk modeling: how to build a pbpk/pd model. CPT: Pharmacometrics and Systems Pharmacology, 5(10):516–531, 2016.

[7] W.-C. Chou and Z. Lin. Machine learning and artificial intelligence in physiologically based pharmacokinetic modeling. Toxicological Sciences, 191(1):1–14, 2023.

[8] US Food and Drug Administration. Physiologically Based Pharmacokinetic Analyses — Format and Content: Guidance for Industry, 2018. FDA Guidance Document.

[9] J. Sheng and T. Zhang. Advancing drug development with “fit-for-purpose” modeling informed approaches. Journal of Pharmacokinetics and Pharmacodynamics, 52(5):52, 2025.

[10] P. Schneider, W. P. Walters, A. T. Plowright, et al. Rethinking drug design in the artificial intelligence era. Nature Reviews Drug Discovery, 19(5):353–364, 2020.

[11] D. Valderrama, A. V. Ponce-Bobadilla, S. Mensing, H. Fröhlich, and S. Stodtmann. Integrating machine learning with pharmacokinetic models: benefits of scientific machine learning in adding neural networks components to existing PK models. CPT: Pharmacometrics and Systems Pharmacology, 13(1):41–53, 2024.

[12] Ali Ghayoor and Hamed Gilzad Kohan. Revolutionizing pharmacokinetics: the dawn of AI-powered analysis. Journal of Pharmacy and Pharmaceutical Sciences, 27:12671, 2024.

[13] A. Vaswani, N. Shazeer, N. Parmar, et al. Attention is all you need. In Advances in Neural Information Processing Systems 30 (NeurIPS 2017), pages 5998–6008, 2017.

[14] Sindhu Tipirneni and Chandan K. Reddy. Self-supervised transformer for sparse and irregularly sampled multivariate clinical time-series. ACM Transactions on Knowledge Discovery from Data, 16(6):1–17, 2022.

[15] T. Chen, S. Kornblith, M. Norouzi, and G. Hinton. A simple framework for contrastive learning of visual representations. In Proceedings of the 37th International Conference on Machine Learning (ICML 2020), pages 1597–1607, 2020.

[16] A. van den Oord, Y. Li, and O. Vinyals. Representation learning with contrastive predictive coding. arXiv preprint arXiv:1807.03748, 2018.

[17] D. A. Boiko, R. MacKnight, B. Kline, and G. Gomes. Autonomous chemical research with large language models. Nature, 624(7992):570–578, 2023.

[18] M. Cranmer, A. Sanchez-Gonzalez, P. Battaglia, et al. Discovering symbolic models from deep learning with inductive biases. In Advances in Neural Information Processing Systems 33 (NeurIPS 2020), pages 17429–17442, 2020.

[19] N. Makke and S. Chawla. Interpretable scientific discovery with symbolic regression: a review. Artificial Intelligence Review, 57:2, 2024.

[20] S. L. Beal and L. B. Sheiner. NONMEM users guide. 1992.

[21] Y. Wang. Derivation of various NONMEM estimation methods. Journal of Pharmacokinetics and Pharmacodynamics, 34(5):575–593, 2007.

[22] R. M. Savic and M. O. Karlsson. Importance of shrinkage in empirical Bayes estimates for diagnostics: problems and solutions. The AAPS Journal, 11(3):558–569, 2009.

[23] M. Rowland, C. Peck, and G. Tucker. Physiologically-based pharmacokinetics in drug development and regulatory science. Annual Review of Pharmacology and Toxicology, 51:45–73, 2011.

[24] E. J. Guest, L. Aarons, J. B. Houston, A. Rostami-Hodjegan, and A. Galetin. Critique of the two-fold measure of prediction success for ratios: application for the assessment of drug-drug interactions. Drug Metabolism and Disposition, 39(2):170–173, 2011.

[25] U.S. Food and Drug Administration. Guidance for Industry: Statistical Approaches to Establishing Bioequivalence, 2001. FDA Guidance Document.

[26] K. Toshimoto. Beyond the basics: A deep dive into parameter estimation for advanced PBPK and QSP models. Drug Metabolism and Pharmacokinetics, 56:101011, 2024.

[27] P. Khosla, P. Teterwak, C. Wang, et al. Supervised contrastive learning. In Advances in Neural Information Processing Systems, volume 33, pages 18661–18673, 2020.

[28] S. Willmann, J. Lippert, M. Sevestre, J. Solodenko, F. Fois, and W. Schmitt. Pk-sim: A physiologically based pharmacokinetic ‘whole-body’ model. Biosilico, 1(4):121–124, 2003.

[29] U.S. Food and Drug Administration. Population Pharmacokinetics: Guidance for Industry, 2022. FDA Guidance Document, February 2022.

[30] C. Lu, C. Lu, R. T. Lange, et al. Towards end-to-end automation of AI research. Nature, 2026.

[31] W. Byon, M. K. Smith, P. Chan, et al. Establishing best practices and guidance in population modeling: an experience with an internal population pharmacokinetic analysis guidance. CPT: Pharmacometrics and Systems Pharmacology, 2(7):e51, 2013.

[32] J. Lu, K. Deng, X. Zhang, G. Liu, and Y. Guan. Neural-ODE for pharmacokinetics modeling and its advantage to alternative machine learning models in predicting new dosing regimens. iScience, 24(7):102804, 2021.

[33] D. S. Bräm, U. Nahum, J. Schropp, M. Clemens, B. Lenz, and M. Pfister. Low-dimensional neural ODEs and their application in pharmacokinetics. Journal of Pharmacokinetics and Pharmacodynamics, 51(2):123–140, 2024.

[34] Y. Cheng, Y. Hu, H. Dong, et al. Exploring transformer model in longitudinal pharmacokinetic/pharmacodynamic analyses and comparing with alternative natural language processing models. Journal of Pharmaceutical Sciences, 113(5):1368–1375, 2024.

[35] A. Das, W. Kong, R. Sen, et al. A decoder-only foundation model for time-series forecasting. In Proceedings of the 41st International Conference on Machine Learning (ICML 2024), 2024.

[36] K. Rasul, A. Ashok, A. R. Williams, et al. Lag-Llama: Towards foundation models for probabilistic time series forecasting. arXiv preprint arXiv:2310.08278, 2023.

[37] M. Moor, O. Banerjee, Z. S. H. Abad, et al. Foundation models for generalist medical artificial intelligence. Nature, 616(7956):259–265, 2023.

[38] N. Terranova, D. Renard, M. H. Shahin, et al. Artificial intelligence for quantitative modeling in drug discovery and development: an Innovation and Quality Consortium perspective. Clinical Pharmacology and Therapeutics, 115(4):658–672, 2024.

[39] K. Stankevičiūtė, J. B. Woillard, and R. W. Peck. Bridging the worlds of pharmacometrics and machine learning. Clinical Pharmacokinetics, 62(11):1551–1565, 2023.

[40] Z. Huang, P. Denti, and H. Mistry. Machine learning and artificial intelligence in PK-PD modeling: fad, friend, or foe? Clinical Pharmacology and Therapeutics, 115(5):652–654, 2024.

[41] A. Talkington, Y. Cao, A. J. Kearsley, and S. K. Lai. Opportunities for machine learning and artificial intelligence in physiologically-based pharmacokinetic modeling. Advanced Drug Delivery Reviews, 227:115716, 2025.

[42] S. Habiballah and B. Reisfeld. Adapting physiologically-based pharmacokinetic models for machine learning applications. Scientific Reports, 13:14934, 2023.

[43] Y. Li, Z. Wang, Y. Li, et al. A combination of machine learning and PBPK modeling approach for pharmacokinetics prediction of small molecules in humans. Pharmaceutical Research, 41(7):1369–1379, 2024.

[44] E. A. Poweleit, A. A. Vinks, and T. Mizuno. Artificial intelligence and machine learning approaches to facilitate therapeutic drug management and model-informed precision dosing. Therapeutic Drug Monitoring, 45(2):143–150, 2023.

[45] A. Janßen, F. C. Bennis, and R. A. A. Mathôot. Adoption of machine learning in pharmacometrics: an overview of recent implementations and their considerations. Pharmaceutics, 14(9):1814, 2022.

[46] M. Ismail, M. Sale, and M. Yu. Development of a genetic algorithm and NONMEM workbench for automating and improving population pharmacokinetic/pharmacodynamic model selection. Journal of Pharmacokinetics and Pharmacodynamics, 49(2):243–256, 2022.

[47] X. Li, M. Sale, J. Craig, K. Nieforth, A. Mazur, and R. R. Bies. Multi-objective optimization in population pharmacokinetic model selection and optimization: application of NSGA-II in pyDarwin. Journal of Pharmacokinetics and Pharmacodynamics, 53(4), 2026.

[48] E. Sibieude, A. Khandelwal, and P. Girard. Population pharmacokinetic model selection assisted by machine learning. Journal of Pharmacokinetics and Pharmacodynamics, 49(2):257– 270, 2022.

[49] Z. Huang. nlmixr2auto: Automated population pharmacokinetic modeling, 2026. CRAN.

[50] Ali Essam Ghareeb, Benjamin Chang, Ludovico Mitchener, Angela Yiu, Caralyn J. Szostkiewicz, Jon M. Laurent, Muhammed T. Razzak, Andrew D. White, Michaela M. Hinks, and Samuel G. Rodriques. A multi-agent system for automating scientific discovery. Nature, 2026. Published online 19 May 2026.

[51] J. Lippert, R. Burghaus, A. Edginton, et al. Open systems pharmacology community—an open access, open source, open science approach to modeling and simulation in pharmaceutical sciences. CPT: Pharmacometrics and Systems Pharmacology, 8(12):878–882, 2019.

[52] S. Willmann, K. Hohn, A. Edginton, M. Sevestre, J. Solodenko, W. Weiss, J. Lippert, and W. Schmitt. Development of a physiology-based whole-body population model for assessing the influence of individual variability on the pharmacokinetics of drugs. Journal of Pharmacokinetics and Pharmacodynamics, 34(3):401–431, 2007.

[53] G. Jocher, A. Chaurasia, and J. Qiu. Ultralytics yolov8, 2023.

[54] R. P. Sheridan. Time-split cross-validation as a method for estimating the goodness of prospective prediction. Journal of Chemical Information and Modeling, 53(4):783–790, 2013.

[55] Z. Wu, B. Ramsundar, E. N. Feinberg, J. Gomes, C. Geniesse, A. S. Pappu, K. Leswing, and V. Pande. Moleculenet: A benchmark for molecular machine learning. Chemical Science, 9(2):513–530, 2018.

[56] J. Grzegorzewski, J. Brandhorst, K. Green, et al. Pk-db: pharmacokinetics database for individualized and stratified computational modeling. Nucleic Acids Research, 49(D1):D1358– D1364, 2021.

[57] F. Fuhrer, D. Galeano, and A. Bender. A deep neural network: mechanistic hybrid model to predict pharmacokinetics in rat. Journal of Computer-Aided Molecular Design, 38(1):7, 2024.

[58] S. M. Kazemi, R. Goel, S. Eghbali, et al. Time2vec: Learning a vector representation of time. arXiv preprint arXiv:1907.05321, 2019.

[59] R. Xiong, Y. Yang, J. He, K. Zheng, et al. On layer normalization in the transformer architecture. In Proceedings of the 37th International Conference on Machine Learning (ICML), pages 10524–10533, 2020.

[60] M. Gibaldi and D. Perrier. Pharmacokinetics. Marcel Dekker, New York, 2nd edition, 1982.

[61] H. Derendorf and S. Schmidt. Rowland and Tozer’s Clinical Pharmacokinetics and Pharmacodynamics: Concepts and Applications. Wolters Kluwer, Philadelphia, 5th edition, 2020.

[62] B. Lakshminarayanan, A. Pritzel, and C. Blundell. Simple and scalable predictive uncertainty estimation using deep ensembles. In Advances in Neural Information Processing Systems (NeurIPS), volume 30, pages 6402–6413, 2017.

[63] M. Cranmer. Interpretable machine learning for science with pysr and symbolicregression.jl. arXiv preprint arXiv:2305.01582, 2023.

[64] P. Virtanen, R. Gommers, T. E. Oliphant, et al. SciPy 1.0: Fundamental algorithms for scientific computing in Python. Nature Methods, 17(3):261–272, 2020.

[65] A. Paszke, S. Gross, F. Massa, A. Lerer, et al. PyTorch: An imperative style, high-performance deep learning library. In Advances in Neural Information Processing Systems (NeurIPS), volume 32, pages 8024–8035, 2019.

